# Augmenting Vascular Disease Diagnosis by Vasculature-aware Unsupervised Learning

**DOI:** 10.1101/2020.02.07.938282

**Authors:** Yong Wang, Mengqi Ji, Shengwei Jiang, Xukang Wang, Jiamin Wu, Feng Duan, Jingtao Fan, Laiqiang Huang, Shaohua Ma, Lu Fang, Qionghai Dai

## Abstract

Vascular diseases are among the leading causes of death and threaten human health worldwide. Imaging examination of vascular pathology with reduced invasiveness is challenging due to the intrinsic vasculature complexity and the non-uniform scattering from bio-tissues. Here, we report VasNet, a vasculature-aware unsupervised learning algorithm that augments pathovascular recognition from small sets of unlabeled fluorescence and digital subtraction angiography (DSA) images. The VasNet adopts the multi-scale fusion strategy with a domain adversarial neural network (DANN) loss function that induces biased pattern reconstruction, by strengthening the features relevant to the retinal vasculature reference while weakening the irrelevant features. VasNet delivers outputs of “Structure + X”, where X refers to multi-dimensional features such as blood flows, the distinguishment of blood dilation and its suspicious counterparts, and the dependence of new pattern emergence on a disease progression, which may assist the discovery of novel diagnostics. Therefore, explainable imaging output from VasNet and other algorithm extensions hold the promise to revolutionize the practice of medical diagnosis, as it improves performance while reduces the cost on human expertise, equipment exquisite and time consumption.

## Introduction

Human organs are infiltrated with blood vessels to supply oxygen and nutrient and eliminate metabolic by-products for living tissue cells^1^. The vascular abnormalities, both structural and dynamic, have strong pathologic indications in a wide range of diseases. Common cerebral vascular diseases, including atherosclerosis, thrombosis, and aneurysm, can cause stroke^2-5^, and many forms of neurological dysfunction and degeneration^6,7^. Inflammatory bowel diseases (IBD), including Crohn’s disease and ulcerative colitis, and bowel microbial infection are found associated with enriched vasculature and microbleeds along the digestive tract^8-10^. Internal bleeding is one major factor leading to mortality after trauma^11^ and hemoptysis^12^. Hypervascular tumor^13^ features enriched angiogenesis. Therefore, probing abnormal vasculature has significant impacts on a range of biomedical examinations.

Vascular imaging with reduced invasiveness causes zero or minimal incision to patients and experimental animals, yet it introduces challenges to the imaging-based diagnosis. Vascular networks are covered by biological structures such as skins and organ tissues, which induce severe light scattering for optical investigations, including fluorescence imaging^14-16^, optical coherence tomography (OCT)^17,18^, and photoacoustic angiography^19-21^. The topologies are imposed with heavy and inhomogeneous noises that disrupt the recognition of pathovascular features. When investigating internal bleeding using X-ray computed tomography (CT) and digital subtraction angiography (DSA)^22-26^, similar pathological features in solid organs, such as vessel bleeding, stricture, aneurysm, and tortuous blood vessels, are difficult to distinguish, due to the artifacts caused by organ motions and the vascular network complexity.

Deconvolution is a relatively common solution to extract information from noise-polluted signals, which usually requires estimating or measuring the point spread function (PSF) of the system based on theoretical models. However, spatially inhomogeneous and individually different characteristics make such non-blind deconvolution solution impossible on biomedical data. Learning-based techniques have recently been utilized to extract high-quality physiological and pathological features and perform recognition across a wide range of scales, including cells^27,28^, histology tissue slides^29^ and deep tissue feature acquired from OCT, CT, and magnetic resonance imaging (MRI)^30,31^. Meanwhile, deep learning has been proven versatile in extracting features from highly scattered patterns through synthetic diffusers^32,33^. Unfortunately, such learning-based descattering methods are supervised, where a large number of labeled and per-pixel registered data are necessary for training neural networks. Moreover, a model trained on one location of the diffuser can hardly be extended to other regions, due to the assumption of spatial-invariant PSF for deconvolution. It has an essential conflict with limited data availability and a large diversity of pathology features. On the other hand, the “black box” operations of deep learning make decisions without explanations. Establishing algorithms to “unbox” learning and generate explainable outputs to render decision-making transparent can accelerate the advances of deep learning in medical diagnosis^34,35^. For example, it might assist the discovery of new diagnostic characteristics, by validating the correlation of initially unexplainable features and the occurrence of a disease.

Herein, we report VasNet, an unsupervised transfer learning technique based on the domain adversarial neural network (DANN)^36^, specialised for vascular feature recognition. It performs vascular-aware domain transfer learning between the widely-accessible retinal vasculature in binary formats as the target domain and the diffusive organ vasculature acquired from fluorescence or X-ray imaging as the source domain. It eliminates the dependence on large datasets of image registration and manual labeling of ground truths (sometimes inaccessible) in supervised methods. The algorithm outputs explainable images with multi-dimensional information of blood flows, including the vascular structure, flow rates, and the pre-screened examination of suspicions. The DANN loss function is embedded in the algorithm to create bias in feature selection to significantly improve vasculature extraction from inhomogeneous backgrounds. In this work, we performed the diagnosis augmentation of thrombosis and internal bleedings and proofed the concept of establishing new diagnostics for ulcerative colitis in animal models.

## Results

### Principle of the vascular imaging augmentation

The concept of augmenting biomedical diagnosis operates by blind vasculature extraction from an unsupervised deep learning algorithm without using labeled data. It is intrinsically an image-to-image translation problem. Recently, some learning-based approaches emerged by using gold standard retinal vascular images to extract explicit vasculature from original fundus images^37,38^. But these models required large datasets of pixel-level aligned image pairs to train the generator network and their performance was limited to retinal vessels. The recent seminal unsupervised image-to-image method, named cycle-consistent generative adversarial networks (CycleGAN)^39^, proved powerful performance on transfer learning between two sets of images, by using cycle consistency loss and adversarial loss to regularize the generator networks and produce realistic images. However, these approaches treat the vascular information and the degradation patterns equally. They were not feasible for eliminating non-uniform noises induced by the heterogeneous diffusibility of skins and organ tissues, uncontrolled organ motions, and non-uniform background illumination.

To generate explainable vascular images using the unsupervised transfer learning technique, we chose a publicly available set of retinal vascular binary images as the target domain reference. (Supplementary Fig. S1) The retinal images presented high contrast topologies of blood vessels against their surroundings with enriched complexity, including the vessel branching, size (diameter) variation, knots, and endings, etc. In the following studies, we implemented VasNet to translate images of cerebral, bowel and internal solid organ vessels, overlayed with heterogeneous noises, into retinal-alike topologies with explainable characteristics.

The principle of diagnosis augmentation is sketched in Fig. 1. A domain-transferred image was consistently generated from an input image containing vascular information and noise overlay by the neural network until the generated image became indistinguishable from explicit binary images of retinal vessels. Instead of presenting an outcome with a binary decision to overlap or validate the human diagnosis, our strategy aimed to augment human diagnosis by presenting the end-users explainable images with enriched information, including the vasculature with improved contrast against the surrounding tissues, the highlights on suspicious regions in colour coding, the probability of disease occurrence, and the trajectory of abnormal blood flows. For example, the coloured vessels represent the longitudinal or transverse flow rates, and the rates in the transverse direction distinguish the syndromes of blood leaking out of the vessels with tortuous vessels and aneurysm. (Fig. 1)

**Fig. 1.**
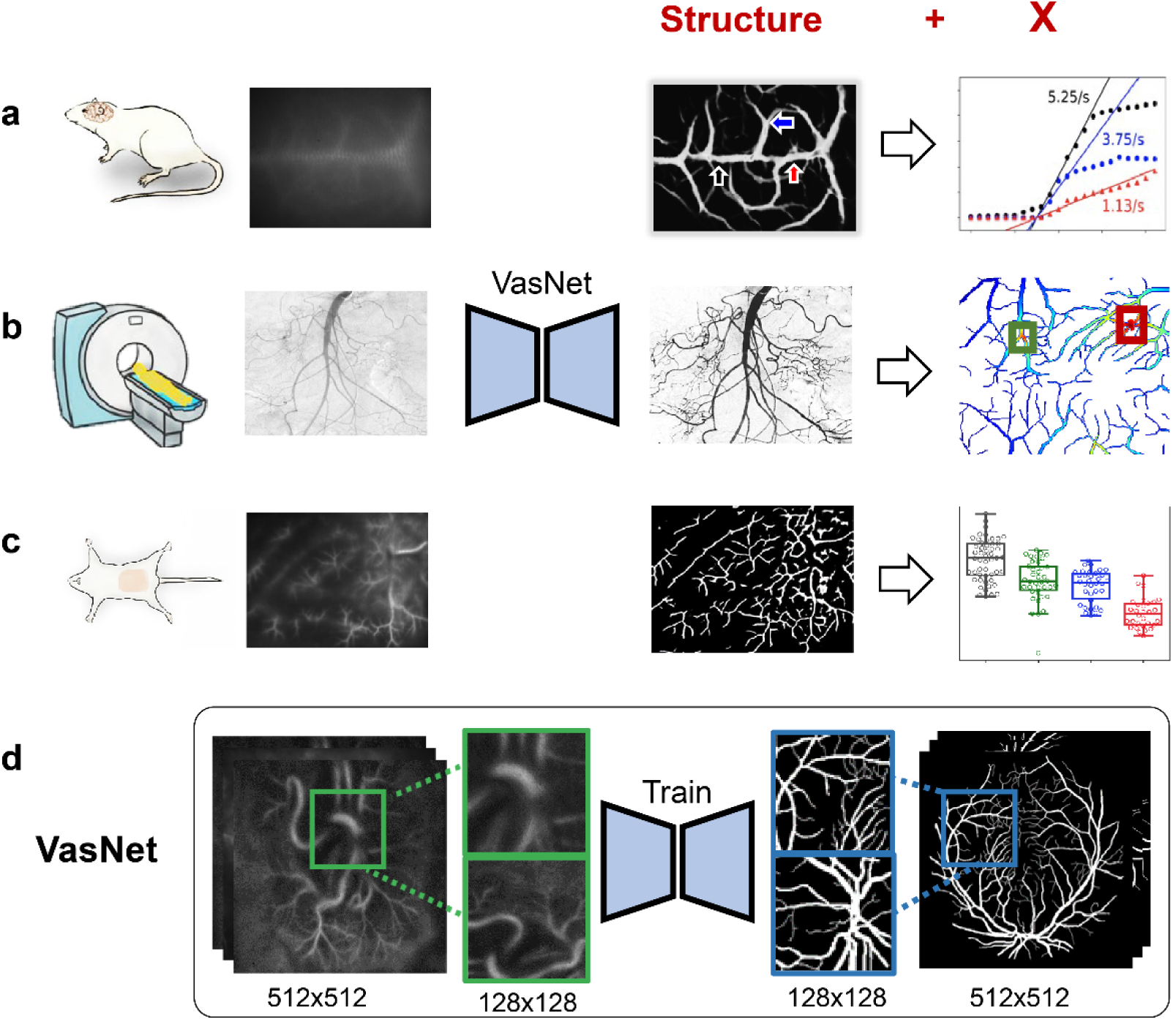
The augmentation principle of vascular disease diagnosis. An unsupervised transfer learning algorithm was established to process and understand raw images corrupted by scattering, aberrations, or non-uniform noises. It extracts the vascular topology, colour-coded the blood flow dynamics, and unveils the spatiotemporal illumination of regions of interest (a), examines the pathological features, and presents the suspicions in contrasting colours (b), and discovers new diagnostic features and suggests the probability of a disease occurrence (c). VasNet learns the image-to-image mapping between two unpaired image domains: the raw vascular observations and the segmentation of the retinal vascular images (d).

Our learning algorithm eliminated the requirement of large datasets of labeled images as ground truths to train neural networks. With this strategy, medical diagnosis is expected to be dramatically improved in many aspects; it reduces the dependence of imaging-based medical diagnosis on high-end equipment; professional users, e.g. doctors, can make faster and more accurate decisions on patients; non-professional users, e.g. patient themselves, medical interns, and general practitioners, can perform pseudo-professional diagnosis. Additionally, it offers a route to discover new diagnosis characteristics, as the unboxing operation generates explainable images, where the subtle morphological changes might be closely correlated with a disease occurrence. In the following sections, we present three pieces of demonstration to prove the versatility of our unsupervised transfer learning potential in different scenarios.

### The VasNet algorithm

The training and testing of biomedical data are often limited by the accessible data volume and their enriched diversity. To recognize vascular features through intact diffusers, such as skins and solid organs that enclose the blood vessels, we eliminated the imposed noises by modeling the non-uniform desattering as an unsupervised mutual information disentanglement problem. Successful descattering was attributed to the adaptation and modification of the unsupervised domain-transfer network, and the selection of the standard dataset of a retinal vascular network as reference for imaging generation. Additionally, we incorporated the multi-scale fusion strategy with a DANN loss function to induce biased pattern reconstruction. Determined by the binary retinal vascular network as the training reference, the noise corrupted images were disentangled into vascular-alike features, also defined as the reference-relevant features, and the noises, defined as the reference-irrelavent features. The VasNet discriminated the two domains after training. Therefore, the algorithm testing performed unblanced selectivity on the vascular-relavent and irrelevant features. As a consequence, the vascular reconstruction was considerably improved and the noise misinterpretation suppressed, compared with the performance of algorithms with non-biased feature selection.

Despite the absence of labeled blood vessels, by utilizing the cycle consistency loss L_*Cycle*_, our domain transfer network (Fig. 2) learned the mapping between two image domains. We fed the segmented retinal vasculature into the VasNet as the target image domain (A domain) and the raw images acquired from fluorescence or X-ray imaging and overlayed with heterogenous noises as the source image domain (B domain). Then, the problem of blind vasculature extraction can be modeled as disentanglement of the vascular structure 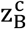 from the degraded observations (real_B). In order to bias the domain transfer by strengthening the vascular-relevant features and weakening the irrelavent features, we incorporated a domain adversarial loss function^40^ to reduce the structural domain shift between the source domain and target domain. The loss penalized the domain gap between the embeddings 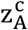 and 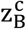, indicating the mutual information of vascular structure. The inference network (coloured in orange in Fig. 2) tended to disregard the vascular-irrelevant features, such as non-uniform background and scattering noises. Furthermore, in order to ease the training process, the network is designed to master only in small scales by restricting the average width of blood vessels in a narrow range. While given the complexity of the vasculature, the algorithm is expected to deal with the blood vessel with different widths simultaneously.

**Fig. 2.**
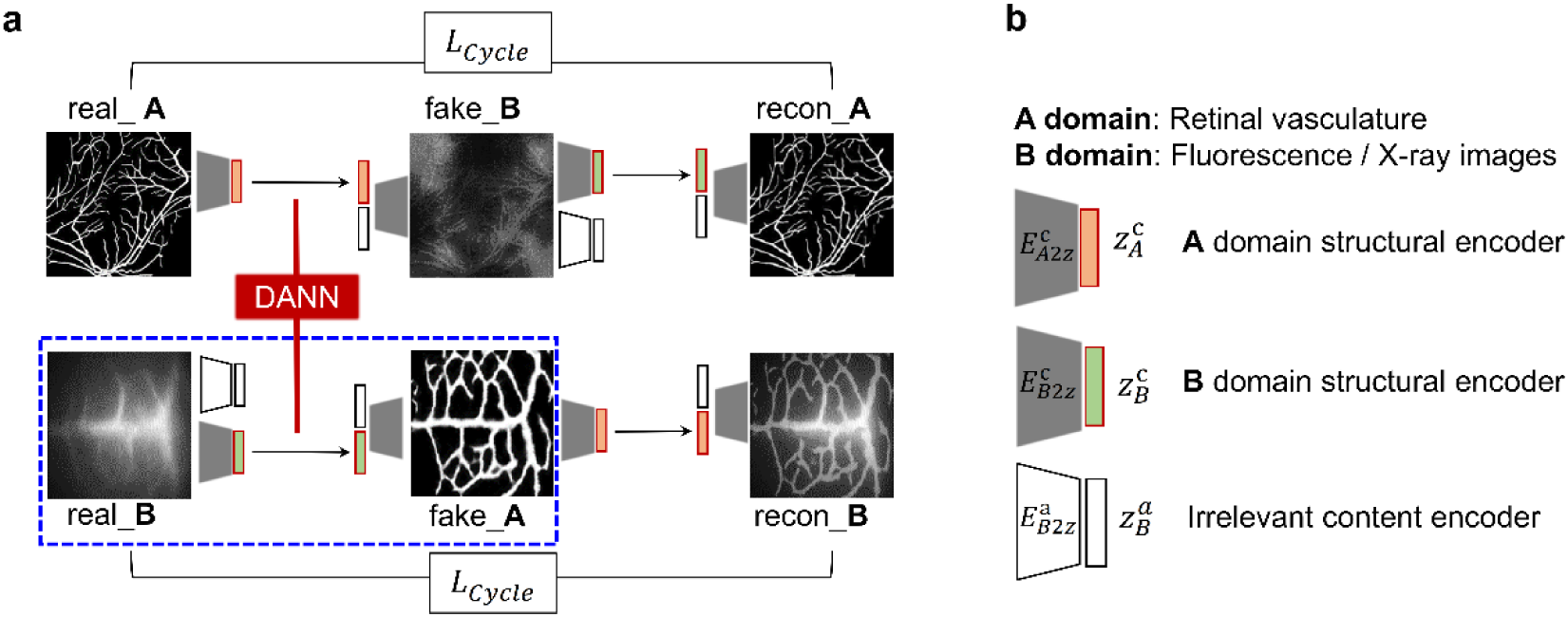
The proposed VasNet solves the heterogeneous descattering problem by extracting the relevant information to the vasculature (domain A) from the scattered observations (domain B). The proposed multi-scale fusion strategy with a domain adversarial neural network (DANN) loss function encourages the emergence of the mutual features, i.e. 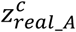 and 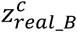.

Therefore, we proposed a coarse-to-fine fusion scheme that merges the predictions of the input pyramid in multiple scales. For each scale, the prediction focuses on the blood vessel with a specific range of width and degree of scattering. Benefited from the vasculature-aware design in the unsupervised domain transfer learning framework, VasNet works regardless of the acquring modalities of the vascular images from the two domains. It delivers the results of “Structure + X”, where X refers to multi-dimensional features such as blood flows, the examined blood dilation and its suspicious counterparts, and the dependence of new structure emergence on a disease progression, which might open a route to assist novel disease diagnostics.

The precise interpretation of vascular network comprising different thicknesses was attributed to the bias in the selection of input topology, created by the DANN loss function in the neural architecture. In this study, as we targeted at the vasculature reconstruction, the algorithm picked the vascular-like features, such as the regional features with high aspect ratios, and tended to disregard the non-vascular features in the reconstruction. This resulted in a significant difference in the output image by using CycleGAN and VasNet (Fig. 3a and 3b). Non-learning approaches were proven even less competitive in processing vascular images. (Supplementary Fig. S2)

**Fig. 3.**
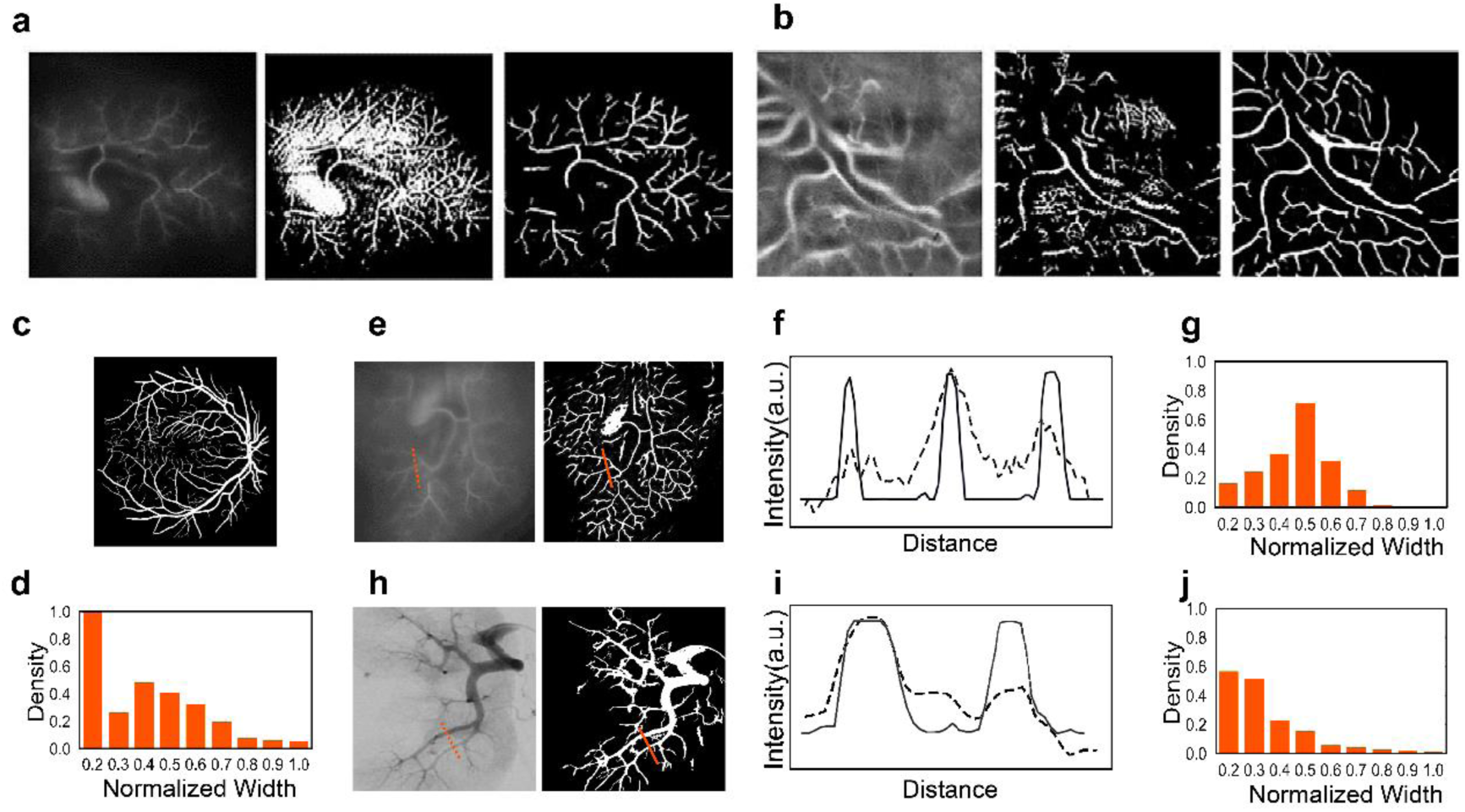
Validation of the VasNet robustness in vasculature reconstruction. (a, b) Comparison of CycleGAN and VasNet in vasculature reconstruction. Each group, from *left* to *right*, has vascular images acquired through intact skins and input into the networks, the reconstructed vascular network from cycleGAN, and the reconstructed network from VasNet. (c - j) The vasculature reconstruction using VasNet is regardless of the vascular topology and distribution. (c, d) A binary image of the retinal vascular network used as the reference (the target domain) for the VasNet training (c) and the count of the vessel width (d). (e - j) The vascular structural reconstruction (e, h), the quantification of regional vessel dimensions (f, i), and the counts of the vessel width (g, j) in the input images and the VasNet output images along the dashed and solid lines in (e) the fluorescent bowel vasculature and (h) the x-ray acquired internal organ vasculature.

A major challenge in imaging-based vascular diagnosis is the loss of structural precision, especially the transverse dimensions of blood flows that indicate the vascular abnormalities, such as blood vessel stricture, dilation, and breakage, etc. The VasNet algorithm compromised the structural loss and recapitulated the topology of the vessels hidden in heterogeneous and complex noises. The versatility of VasNet on different imaging techniques with variations in vascular structures was validated by comparison. The reference retinal vascular image had enriched variation in vessel topology and size, but the distribution of vessel dimensions in the testing images was distinctive with the reference images, regardless of the non-uniform noise magnitude and distribution overlayed on the patterns. (Fig. 3c – 3j)

### The DANN biased feature reconstruction

Prior to proceeding to the feature recognition from diffusive patterns in experimental models and humans, we first tested the performance of the DANN loss function on biased feature selection on a simulation system. A digital micromirror device (DMD) projected letter patterns on a non-transparent polystyrene board or a piece of 3 mm-thick chicken breast slice, which were acquired by a camera from the other side of the diffuser. (Supplementary Fig. S3) The diffuser was placed on a translation stage to create the inhomogeneous diffusion that mimicked the *in vivo* conditions. The acquired patterns and the reconstructed features by using the reported deep learning algorithms (‘Li et al. 2019, the reference 32’, with fixed diffusivity) and our algorithm (‘ours’) are presented in Supplementary Information. The comparison showed that our DANN algorithm gained both improved feature extraction ability from diffused speckles and higher versatility in overcoming the disturbance caused by the changes in diffusivity. (Supplementary Fig. S4 – S6)

### Augmenting fluorescence imaging for the diagnosis of cerebral vascular disease

The Indocyanine green (ICG) is an FDA approved dye molecule for clinical uses. It has emission in the near-infrared window (the peak emission at ∼ 820 nm). The specific binding of ICG to lipoproteins in the bloodstreams prolongs the circulation lifetime of ICG in blood^14^. Though the light scattering of skins weakens with increased wavelength, the acquired images in our experiments in the NIR-I region (800 - 900 nm) were proven highly disrupted for feature recognition at zero or minimal tissue incisions. Herein, the unlabeled diffused images were input into the transfer learning network and translated into explainable images with both precision vascular structures and spatial flow dynamics in the mouse cerebrum.

The cerebral vasculature was covered by a translucent skull and an intact scalp skin (about 0.6 mm thick). It had a stem vein along the superior longitudinal sinus, with branched vessels distributed on the two sides. A stroke occurs when the blood supply is occluded. Thrombosis is a common type of occlusion resulted from the blood clot formation inside vessels. A mouse model of thrombosis was created by intravenously injecting Rose Bengal and exposing a region of the cerebrum to a 532 nm laser light (0.8 W) for 4 min through the intact scalp skin and the skull^41^. Ischemic lesion was induced, obtaining the thrombosis model for diagnostic tests in the following study. (Supplementary Fig. 7a)

The ICG was first dissolved in phosphate-buffered saline (PBS). Then, a BALB/c mouse was intravenously injected ICG at a dosage of 8.0 mg/kg body weight (bw) at once. The dye molecules rapidly circulated to the cerebrum within a few seconds after the intravenous injection. A 785nm laser beam was projected on the mouse head with the topical hair removed. (Supplementary Fig. 7b) The fluorescence from ICG in circulation that illuminated the cerebral vessels were recorded immediately after ICG injection at a frame rate of 25 Hz through an 810 – 890 nm bandpass filter. The bright-field images of the vascular structure were acquired as the reference after removing the scalp and the skull. The healthy mice were imaged in the same procedures as the diseased ones for control.

The acquired image sequences contained both the spatial and time-dependent illumination of ICG circulating in the blood, from the variation of which could the blood flow rates be derived. Owing to the light scattering through the scalp and the skull, only diffused vascular topology of healthy and diseased mice could be acquired. (Fig. 4a and 4c) We applied VasNet to eliminate the irrelevant information from the vasculature, including noises and non-vascular topologies, and extracted precise vascular structures from the relevant signals. 10 retinal vascular images of 512 × 512 size were fed to VasNet as the target domain (the real domain A in Fig. 2), and 10 unlabeled images from fluorescence acquisition with a size of 1024 × 1024 as the source domain (the real domain B in Fig. 2). Note that the augmented training data is sampled from the large images with the patch size of 128 × 128. The algorithm-generated images exhibited high-contrast vascular features (Fig. 4b and 4d) that were validated by the patterns obtained after removing the scalp (Supplementary Fig. S8). The global cerebral vasculature was constructed from the illumination sequence.

**Fig. 4.**
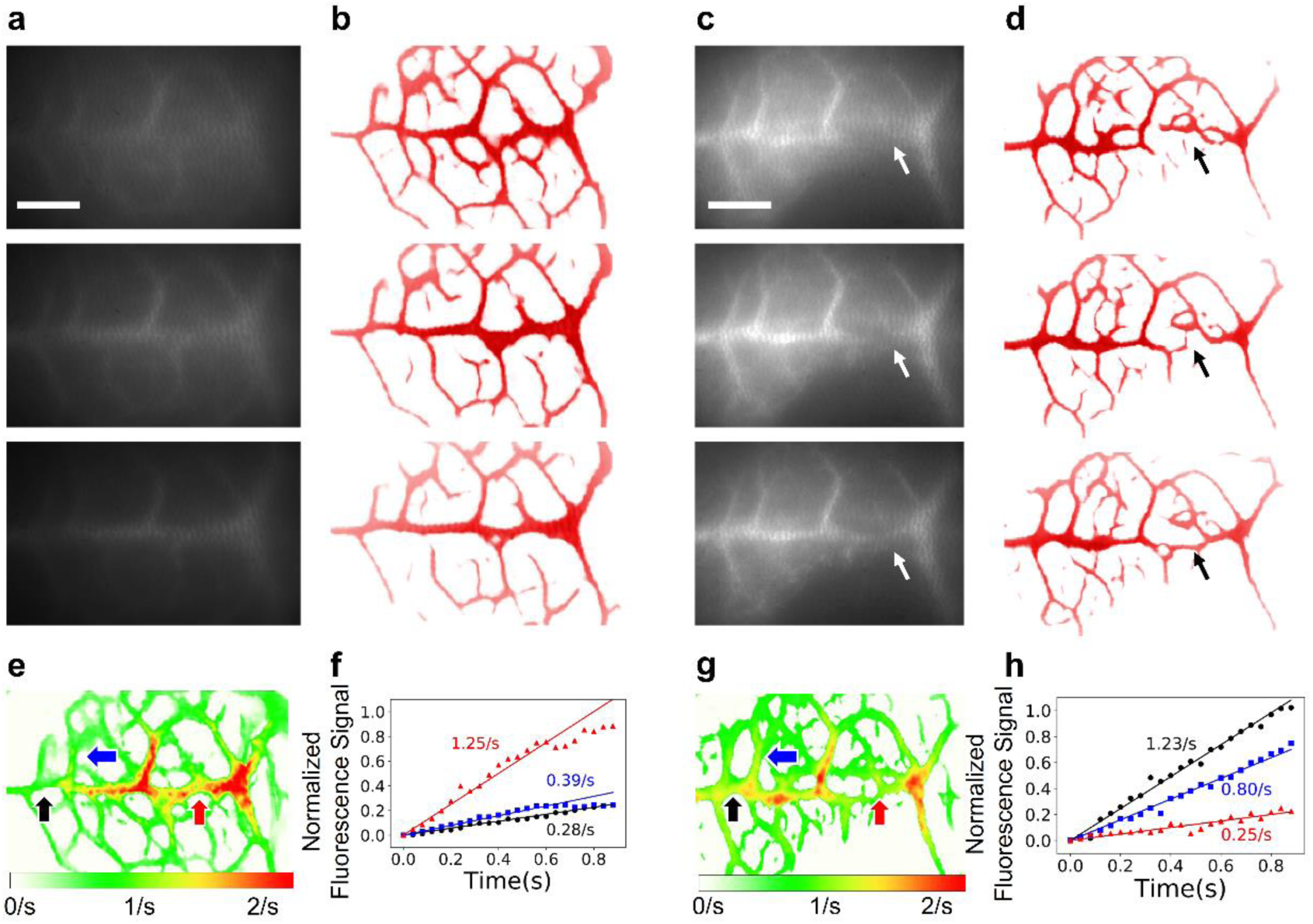
Fluorescence imaging and interpretation of the cerebral vasculature using the unsupervised transfer learning algorithm. The vascular network was perfused with ICG and imaged in the near-infrared window (excitation at 785 nm and emission at 810 - 890 nm). (a - d) Acquired images of ICG illuminated vessels (a, c) and the VasNet output images (b, d) in the cerebrum of a healthy mouse (a, b) and a mouse with thrombosis (c, d). (e - h) The derived blood flow rates from the time-dependent illumination of blood vessels in a healthy cerebrum (e, f) and a diseased cerebrum (g, h). The time-dependent intensity variation on multiple locations indicated with coloured arrows were plotted in (f, h). Scale bars in all panels: 2 mm.

The blood flow rates throughout the network were derived from the illumination sequence, and encoded in colours, based on the principle of the continuum of mass for incompressible fluids, i.e. the blood. (Fig. 4e) The colour-coding shows predominantly drops in regional flow rates resulted from blood occlusion in the thrombosis models, when compared with their healthy counterparts. The flow rates and their variations in different parts of the vascular network were found consistent in at least 3 healthy and 3 diseased mice. (Fig. 4e – 4h, and Supplementary Fig. S9) It proved that our algorithm was not only reliable in probing the structural abnormalities, but also the abruption in flow rates with high spatial signatures in the cerebrum. The structural-dynamic dual-check assured augmented higher diagnosis efficiency and accuracy in the non-invasive examination.

### Augmenting DSA imaging for the diagnosis of internal vascular disease

Digital subtraction angiography (DSA) is among the most important examinations in the diagnosis and treatment of internal bleeding and hypervascular tumors via intervention embolization^25,26^. However, patient motions, respiration, and the skeletons create severe artifacts and impose noises onto the vascular features, which may result in a delayed diagnosis or a high rate of misdiagnosis. For example, bleeding, tortuous vessels, and aneurysm are all characterized by dilation of the radiocontrast agents, and the overlay noises lead to indistinguishment between the pathological vascular features and their normal counterparts.

Though the dilation of the radiocontrast agent (iodixanol) was observed from the DSA images (Fig. 5a and 5c), the artifacts caused by respiration induced organ motions reduced the recognition of suspicious features or features hidden within the complex vascular background. Furthermore, the intervention embolization required a high-rate, or nearly real-time, diagnosis of diseased features with precision spatial resolution, as it was operated under the real-time guidance of DSA imaging.

**Fig. 5.**
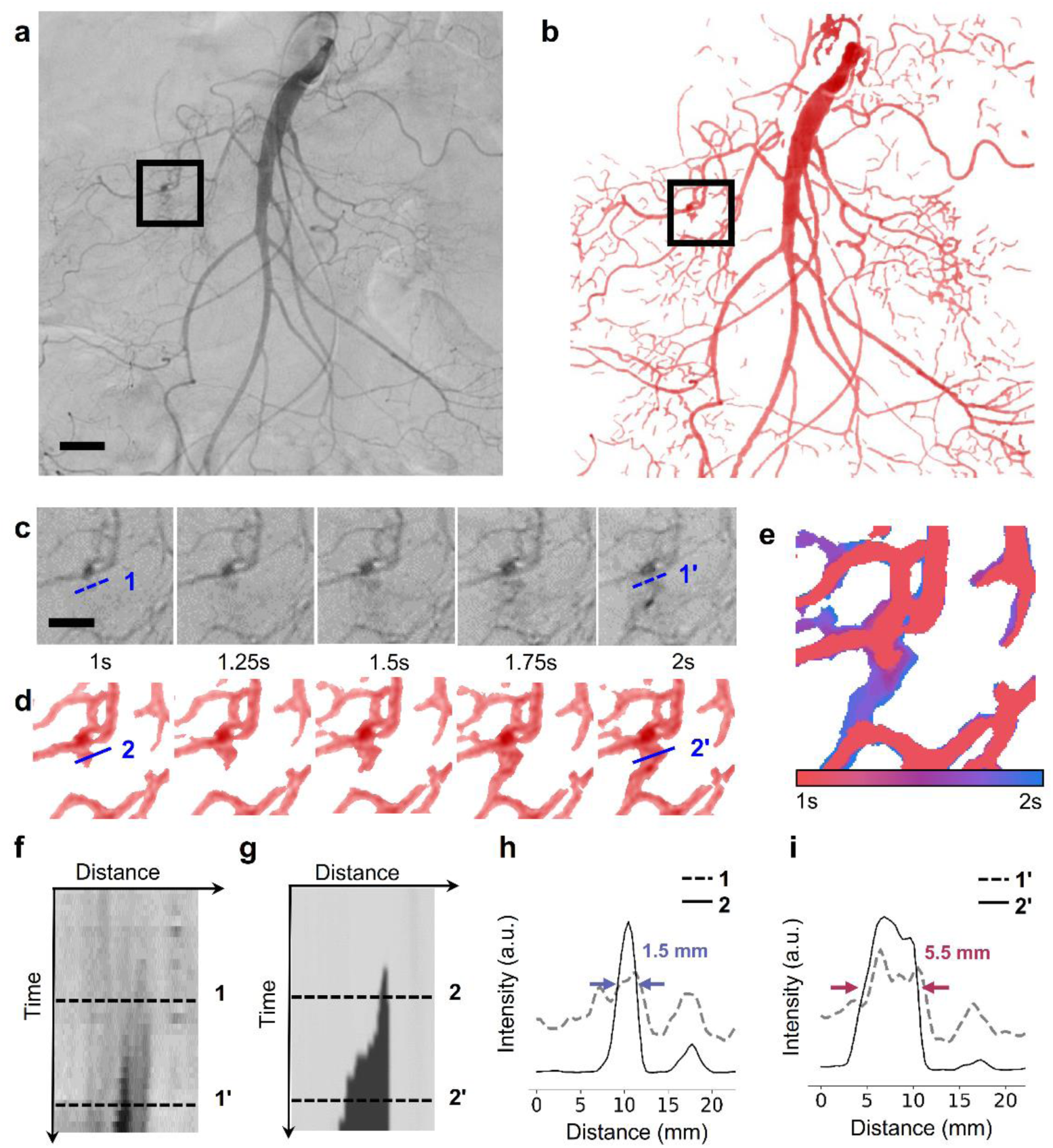
The VasNet interpretation of time-lapse DSA vascular images. (a) A DSA image of the internal vasculature. The images were acquired with a radiocontrast agent (iodixanol) to highlight the blood circulation. (b) The VasNet output image with the vasculature reconstructed and the bleeding features augmented. (c, d) Zoom-in time-lapse view of the regions of interest (ROIs) (framed) in (a, b). The dashed (lines 1 and 1’) and solid lines (lines 2 and 2’) in the image sequence indicate the dilation. (e) The dilation trajectory of iodixanol at the bleeding site, derived from the VasNet output sequences in (d). The colour-coding shows the dynamics of dilation. (f, g) The time-dependent intensities of iodixanol along the dashed and solid lines in (c, d). (h, i) Quantification of the iodixanol intensities along the dashed and solid lines in (c, d) at two time-points. Scale bars: (a) 20 mm, (c) 10 mm.

The limitation was overcome by generating explainable images from the VasNet algorithm. (Fig. 5b, 5d, 5e) The global vascular network was reconstructed with significantly improved accuracy and contrast against the background noise. The ROIs framed in black was interpreted with colour-coding exhibiting the dilation in the recording. The dilation recognition in the raw input and the output images were compared. (Fig. 5f – 5i) The intensity contrast of the circulating agent in the dilation track, contributing positively to the diagnosis ability, was less distinctive in the DSA raw images than in the learning output images. The feature augmentation was performed in 40 groups of DSA image sequences, each acquired from a patient individual. Three examples are shown in Fig. 5 and Supplementary Fig. S10.

The VasNet interpretation also augmented diagnosis in single static images. Examining the bleeding vessel, it is featured with a narrow neck connecting to an expanded domain, corresponding to the dilation of blood and the radiocontrast agent. However, the dilation was not clearly visible in the non-augmented images due to the elusive boundaries. The narrow-neck connected expansion was not observed in the branched or coiled vessels or aneurysm.

VasNet enhanced the vascular structure against the non-uniform degradation, and succeeded to deliver outputs with “Structure + X” in DSA modality, where X helps with the distinguishment of blood dilation. To validate the effectiveness of augmenting bleeding diagonosis quantitatively, we particularly trained a basic classifier to determine whether the bleeding occurred or not, tested on the augmented vasculature of the suspicious patches. More specifically, the data from 40 patients were collected, where half of them were adopted for training, and the rest for testing. Note that the markers of bleeding region were provided by clinicians in the Chinese PLA general hospital, and used as ground truth data for accuracy calculation.

The correctly classified positive (red) and negative (blue) samples are depicted in Fig. 6a and 6b. By interpreting the bleeding recognition results, we conclude that the classifier prefers to group the augmented samples with narrow-neck connected expansion together as positive predictions. For the case exampled in Fig. 6c that counted the bleeding and non-bleeding features of 20 patients, the bleeding diagnosis performance by the neural network, quantitatived by the area under the ROC curve (AUC), reached 97%. The classification accuracy reached 88% when the cut-off probability was 0.4.

**Fig. 6.**
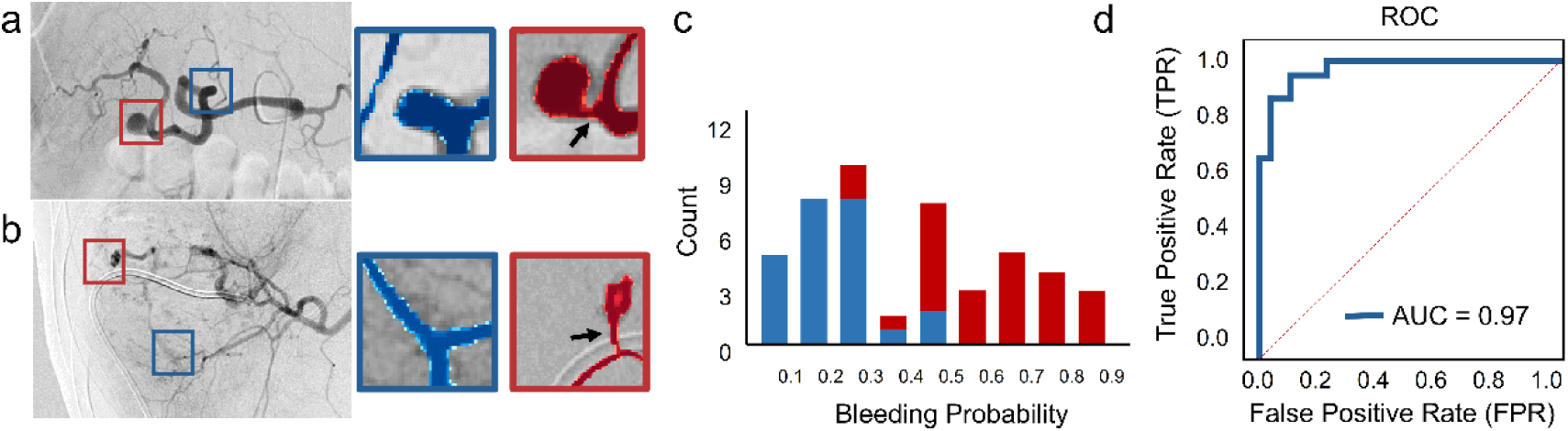
The diagnosis augmentation of bleeding in static DSA images. (a, b) The distinctive feature recognition to distinguish aneurysm (blue in a), tortuous vessel (blue in b) and bleeding (red in a and b). The red vessels were featured with thin necks and expanded domains. (c) The counts of bleeding (red) and non-bleeding suspicions (blue) in the DSA images (20 patient samples in total) with each count showing the bleeding probability. The bleeding and non-bleeding features were distinctive by interpreting their bleeding probability. (d) The receiver operating characteristic (ROC) curve of bleeding counts.

### Bowel vascular network

Inflammatory bowel diseases (IBD), mainly comprising ulcerative colitis (UC) and Crohn’s Disease, have mucosal inflammation and are featured with an increased density of abnormal vessels, such as strictures, ulcers, bleeding along the tract, as well as mucosal angiogenesis^9,10^. Mouse colitis models were established by watering BALB/c mice with dextran sulfate sodium (DSS, 5 wt% in drinking water). DSS carries a highly negative charge contributed by sulfate groups, is toxic to the colonic epithelia, and induces erosions that ultimately compromise barrier integrity, resulting in increased colonic epithelial permeability^42^. In optimal conditions, disease induction occurred within 3 to 7 days following DSS administration, and appeared as severe colonic bleeding. It mimicked the superficial inflammation seen in ulcerative colitis, which was found featuring enriched angiogenesis in the mucosa, but nothing reported on the bowel outer surface^43^.

Likewise, we used ICG to enhance the probing of the bowel vasculature. The bowel vasculature was covered by the abdominal skin (about 0.3 mm thick), which scattered the emission from the circulating ICG in the vessels that appeared on the surface of the tract. A mouse was fixed on the imaging table, with the hair-removed abdomen facing the camera fixed on the top. The abdominal vasculature was illuminated by an expanded 785 nm laser beam, with an illumination area and field-of-view larger than 9 cm^2^. Both healthy and DSS treated mice were imaged in the same manner, by recording their spatiotemporal illumination of the bowel vasculature at high frequency (25 Hz), started immediately after the ICG injection from the tail vein. The images were acquired at days 2, 4 and 6 after the DSS administration. The healthy mice were imaged as the control, named as day 0 in the counting. The ground truths of the health conditions were accessed by cutting off the abdominal skin and the peritoneum with iris scissors and presenting the vascular features to the camera. (Fig. 7)

**Fig. 7.**
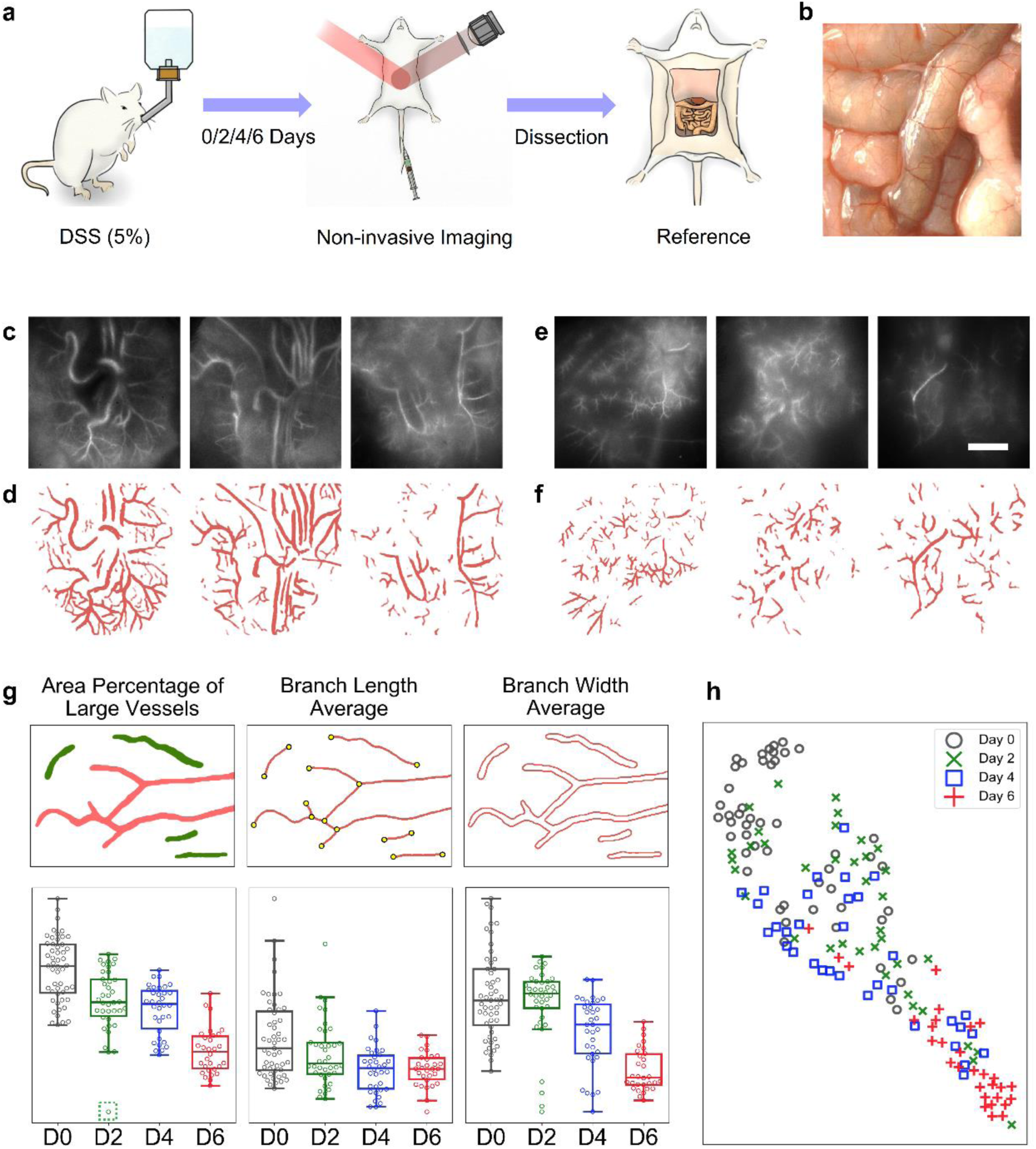
Fluorescence imaging and interpretation of the mice bowel vasculature to diagnose colitis. (a) Steps of diagnostic testing, including creating the colitis models by watering mice with DSS, non-invasive imaging of the bowel vasculature, and access to ground truths of the vasculature by sectioning off the abdominal skins. (b) Bright-field image of the bowel vasculature of an experimental mouse. (c - f) Acquired fluorescence images (c, e) and interpreted images (d, f) of the bowel vasculature of healthy mice (c, d) and mice with colitis (e, f). (g) The multi-style interpretation and counts of bowel vasculature, including the area percentage of large vessels, the average branch length, and the average branch width. (h) The scatter diagram of blood vessels acquired from 22 mice based on quantitative evaluation of the area percentage of large vessels, the average length and the average width of the vessels and their branches. We used the T-distributed Stochastic Neighbor Embedding (t-SNE) algorithm to embed high-dimensional parameters in 2-dimensional space for visualization. Scale bar: 5 mm.

The explicit vascular topology extracted from the blurry raw images was interpreted with various diversification methods, including the large vessel fraction, and the vessel length and width. (Fig. 7g, 7h) Fig. 7g shows the statistics of vessel diversification of mice at days 0, 2, 4, 6 after being administrated with DSS, by measuring the vessel connectivity, the average length and the average width of the reconstructed vessels. The connectivity was quantified by evaluating the fraction of connected vessels greater than a particular value in area (150 pixels in this case), indexed as ‘area percentage of large vessels’. The percentage was calculated as the proportion of the sum pixel value of large vessels over all vessels in each image. The average lengths and the average widths were quantified by Fiji^44^. The quantification was presented in the corresponding boxplots in Figure 3g. For better visualization of the multiple indices, we performed the t-SNE algorithm^45^ to present the diversification in two dimensions.

The statistics show the distinct distribution of mice groups at different health conditions, as suggested by their disease progression, indexed by the number of days after the DSS administration. (Fig. 7h) However, the distinction was not always the same, as the separation by calculating the area fraction of large vessels is the most significant, but the least significant in their width statistics. The outlier at day 2 in the connectivity plot, indicated by a dashed bracket, was suggested to be heavily diseased. It was verified by the skinless observation of the bowel conditions. (Supplementary Fig. S11) This study extended the promise of our unsupervised transfer learning technique in discovering new diagnostic characteristics, which is beyond the scope of the reported artificial intelligence in medical diagnosis.

## Discussion

In this work, we reported a new class of contribution that the deep learning technology might provide to humans, by showing the power of machine intelligence in augmenting diagnosis of vascular diseases. We established VasNet, an unsupervised transfer learning network to overcome the challenges raised by the limited data accessibility in biomedical imaging, the large diversity of pathological features, and the labor-intensive image registration and ground-truth labeling for training a neural network. In the algorithm, we embedded a multi-scale domain adversarial training scheme that simplifies the disentanglement between the vascular structure and the modality-specific noise representation. It was proven versatile for the interpretation of images acquired from different modalities and carrying structures with varied dimensions, size distributions, and heterogeneous diffusion levels. Rather than providing binary decision-making outcome, VesNet unboxed the feasibility of learning on disease diagnosis, and generated explainable images with multi-dimensional information, including vascular topology, flow rates, prescreened suspicions and statistics of blood vessel features. Therefore, the acceptance of artificial intelligence (AI) decision-making is boosted. The doctors could either accept a binary dicision or make decision based on the enriched information output from the neural network. Consequently, it significantly decreased the rate of misdiagnosis, reduced the diagnosis cost as it required less time to decide and became less dependent on high-end equipment.

We also proposed the idea of using deep learning techniques to discover novel diagnostic characteristics. Unknown characteristics are usually hidden under heterogeneous noises, as the vascular patterns shown in Figure 7c and 8d. Though the distinction was observable from the raw images, it requires clear and amplified differences in the pattern evolvement with the disease progress to distinguish the features in healthy, pseudo-healthy and diseased conditions. Our algorithm-dependent examination amplified the elicit distinction, and therefore, validated the diagnosis of colitis by their vasculature appearance on the bowel exterior surfaces. The colitis progresses with angiogenesis, which initiates at the early stage of disease occurrence.

The technology could be extended to the diagnosis of other types of diseases, depending on the feature classification and diversification. For example, lymphatic disorders, such as lymphedema (LED), by lymphoscintigraphy (LSG), but this technique is limited in its ability to identify pathology and guide therapy. Our AI-based medical imaging augmentation might not only assist the rapid diagnosis of LED, but is also promising to guide therapeutic intervention^46,47^.

Our diagnosis augmentation strategy offers a route for non-experienced users to make rational decisions on symptoms on either experimental animals or patients. The AI augmented diagnosis takes over the loads on doctors and experimentalists by reducing the dependence on personnel experience, equipment qualities, and repetitive labor intensive practice and confirmation. With the advent of new diagnosis technologies, such as portable, easily affordable, and family-equipped diagnostic instruments, patients could make preliminary judgement on their health conditions. By all these manner, the deep learning technology holds the promise to alter the way of healthcare and hospital management.

## Methods

All the animal studies and usage of clinical data were conducted under the ethics regulation of Tsinghua University and the PLA General Hospital, China.

### Fluorescence imaging setup

In both imaging experiments through the scalp and abdomen, the excitation light was provided by a 785 nm laser source (Changchun New Industries) coupled to an 8 mm collimator. Light was expanded and adjusted by an iris to illuminate the entire mouse head or abdomen area. The excitation power density at the imaging plane was about 29 mW·cm-2 in our experiments. The emitted fluorescence was filtered by an 810 - 890 nm bandpass filter (Thorlabs, FBH850-40) and captured by a CMOS camera (Blackfly, BFS-U3-120S4M-CS) through a lens with a focal length of 25 mm. The camera was set to expose continuously in a frame rate of 25 f.p.s. to the fluorescence illumination process.

### Animal preparation

All animal experiments were performed in accordance with the National Institutes of Health Guide for the Care and Use of Laboratory Animals and approved by the Institutional Animal Care and Use Committee of Tsinghua University, China. BALB/c mice were purchased from Guangdong Medical Laboratory Animal Center. Adult BALB/c mice (male, 4-6 weeks old) were housed at 22 ± 2 °C with half-to-half light-dark cycle and were used with randomization or blinding. The mice were intraperitoneally injected at a dosage of 240 mg/kg body weight (bw) 1.25% avertin at once. Before imaging, the hair on the heads and abdomens of the mice was removed by using the human depilatory cream.

### Cerebral thrombosis mice model

BALB/c mice were anesthetized by isoflurane, and the hair over the scalp was removed. Then the mouse was intravenously injected Rose bengal at a dosage of 10 mg/kg body weight (bw) at once. The mouse head was exposed under a 532 nm laser for 5 min and the radius of the spot was about 2.5 mm. Rose Bengal irradiated with green excitation light generates the production of reactive oxygen species, which subsequently activates tissue factor, an initiator of the coagulation cascade. The induction of the coagulation cascade produces an ischemic lesion that is pathologically relevant to clinical stroke^48^. After 1 day, the mice were used to conduct the cerebral vasculature imaging.

### Ulcerative colitis mice model

BALB/c mice were randomly assigned to the experimental or control groups (n = 5 - 6). For the experimental group, 5 wt% dextran sodium sulfate (DSS) in double-distilled water (DDW) was provided as drinking water for 2, 4 or 6 days. For the control group, DDW was always used as the drinking water. The treated and untreated mice were imaged on the abdomen on the same day.

### Vasculature imaging

A BALB/c mouse with the hair removed on the head or abdomen was anesthetized by isoflurane and fixed under the camera and lens. The range of interest(head or abdomen) was exposed under a 785 nm laser source. Then, the mouse was intravenously injected Indocyanine green(ICG) at a dosage of 8 mg/kg body weight (bw) at once. The ICG molecules transport rapidly with the circulation, and the vasculature became fluorescent almost instantaneously after the intravenous injection. Camera acquisition of the fluorescence images started right before the intravenous injection to record the illumination process.

### Preparation for the DSA imaging dataset

We collected our angiography data set of 35 patients from the Chinese PLA General Hospital, Beijing, China. For each patient, 3-6 images during the dynamic process containing the vascular information were annotated the real bleeding point region(true label) and 2-3 normal yet suspicious regions(false label). To perform the bleeding detection task, DSA images of 15 patients were used for training and images of 20 patients were used for testing.

### Learning algorithm

To perform the training and testing of biomedical data with limited volume and enriched diversity, we innovated the unsupervised neural network to emphasize feature reconstruction for vascular disease diagnosis, denoted as VasNet. The success was attributed to the adaptation and modification of the domain-transfer network and selection of the standard dataset of a retinal vascular network as the reference for imaging generation. The incorporation of a multiscale fusion strategy with DANN loss function creates a biased reconstruction on vascular-relevant patterns. Consecutively, VasNet significantly improved vasculature extraction and reduced the misinterpretation of the heterogeneous noises.

### Network structure

As shown in Figure2, VasNet decouples the structural information 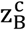 and 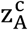 from the source domain (B) and destination domain (A) by the encoders 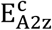 and 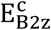 and the irrelevant content 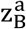 by the encoder 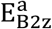. The destination image domain is expected to contain the vascular structure of interest. Besides the vasculature content, the source image domain suffers from various types of degradations, such as non-uniform background and non-uniform degree of scattering, decoupled by the decoder 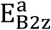. In order to effectively decouple the vasculature content, i.e. the small gap between the distributions of 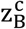 and 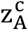., we propose to penalize the domain invariance between 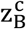 and 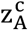 by the domain adversarial loss proposed in the DANN^36^. In addition, since the DANN loss may lead to a small gap between two feature distributions, none of which is fixed, the oscillation in the training process commonly existed, as mentioned by Tzeng et al.^49^. To deal with it, the domain adversarial loss adopted a cross-entropy loss function against random guess result. Such domain adversarial loss encourages the emergence of indiscriminative structural information from both source and destination domains. Since the mutual information between 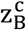 and 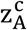 is the vasculature, the domain adversarial straining scheme ensures the success of vasculature-aware unsupervised learning algorithm for the case of non-uniform background and scattering.

Considering the existence of various width of the blood vessels, we further utilize a coarse-to-fine scheme by fusing the predictions of the input pyramid with different scales for both wide and narrow vessels. Another way to tackle the problem is training with multiple networks for each input scale. Unfortunately, we find that it significantly slows down the training process and dramatically requires large data volume. In contrast, taking advantage of the fact that various scales of the vasculature share the similar topology and width distribution, our multiscale fusion strategy leads to considerably complete and accurate vasculature reconstruction.

### Training data preparation

In order to bias the vasculature-aware pattern reconstruction, we explicitly collect the destination domain images based on the desired structure features. The publicly available retinal vasculature dataset CHASE_DB1^50^ and DRIVE^51^ with manual segmentation is selected as the destination domain image. Since the vascular structure of different scale shares similar topology, the algorithm merely requires for inputting small patches (128×128) cropped from the raw large images. Similarly, the source domain images, which are from different modalities and have various degrees of scattering, are also cropped into small patches. The criteria to determine the patch scale is to make the histogram of average vascular branches length from the two domains resemble each other.

## Acknowledgments

The work was supported by the national natural science foundation of China (Grant Number: 61722209, 61971255), the fund from the Shenzhen Science and Technology Innovation Committee (Grant Number: KQJSCX20180327143623167), and the fund from Shenzhen Development and Reform Commission Subject Construction Project [2017]143.

## Author Contributions

L.F., S.M., and Q.D. conceived this project. S.M., L.F., Y.W., S.J., and M.J. designed the experiments. Y.W. and S.J. performed animal experiments. M.J. innovated and implemented the algorithm. M.J., X.W., and J.F. tested the algorithm and performed the pre- and post-processing of the data. F.D. contributed the X-ray and DSA data under the ethical approval from the PLA general hospital. Q.D. oversaw the experimental design and progress. S.M., L.F., M.J., Y.W., and J.W. prepared the manuscript. All co-authors reviewed the paper.

