## Supplemental Material for "Augmenting Vascular Disease Diagnosis by Vasculature-aware Unsupervised Learning"

### 1. Training set used in domain adaptation

The blurry images captured in our animal experiment and the DSA images are to be transformed into the target domain, i.e., the domain of explicit binary images. Ground truths of the above biomedical images were not easily accessible, so we used the standard retinal vascular images in binary formats, acquired from the STARE project as the reference to conduct our domain adaptation tasks.

Examples of the retinal vascular images are shown in Supplementary Fig. S1.

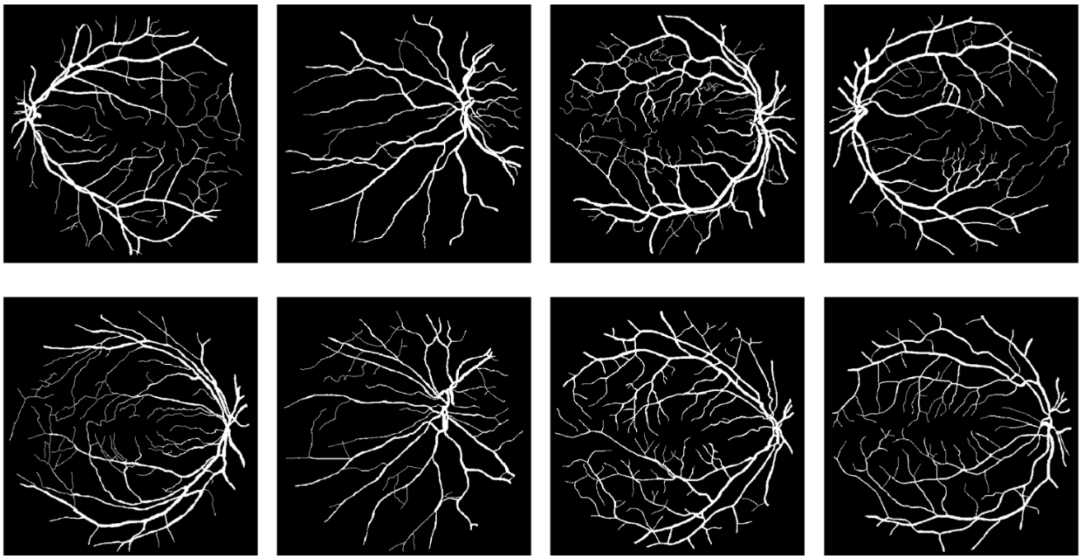

**Fig. S1** Examples of the retinal vascular images used in the VasNet training processes.

### 2. Performance comparison of VasNet against the existing descattering techniques

To test the competence of the VasNet algorithm in tackling various descattering problems, especially eliminating heterogenous noise backgrounds from biomedical vascular images, we compared the performance of VasNet with other computational techniques, inclduing the adaptive thresholding, the vessel analysis function in Fiji, and the deconvolution processing. Our VasNet was proven advantageous in eliminating noises and extract vascular feature from the non-uniform background levels.

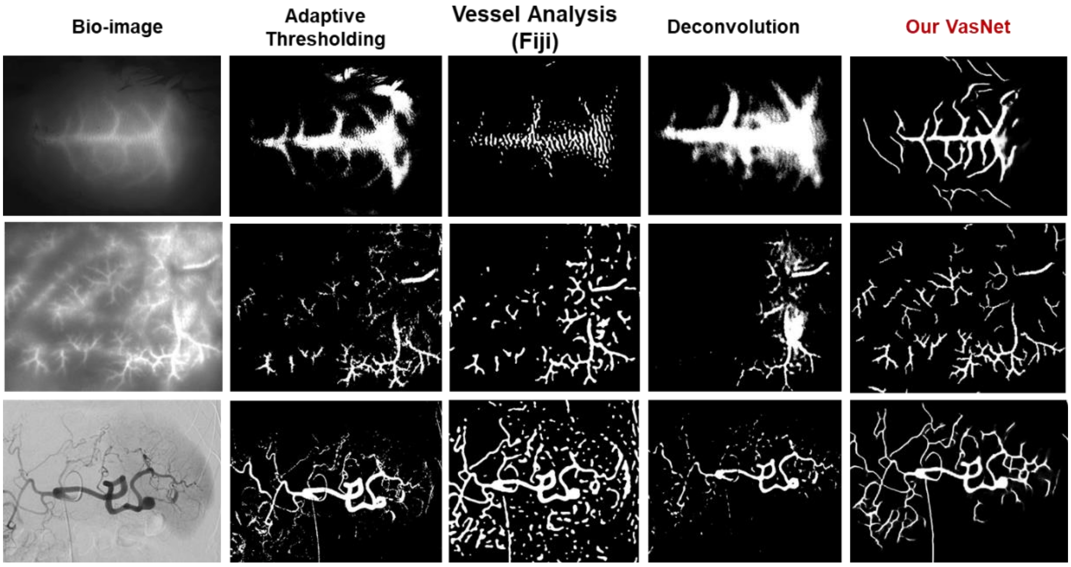


**Fig. S2** Performance comparison of vasculature reconstruction using different techniques. First column: raw biomedical images. Second column: results by applying adaptive thresholding algorithms. Third column: results from the vessel analysis in Fiji. Fourth column: results by applying the deconvolution algorithm. Fifth column: vasculature generated by VasNet.

#### 3. *In vitro* experiment using synthetic diffusers for principle validation of domain adaptation and biased feature reconstruction

Extracting recognizable features from scattered optical patterns remains a common challenge in non-invasive biological and medical examination, as biological tissues, including skins, are high scattering media. To test the de-scattering ability of DANN, we assembled an optical system with a polystyrene (PS) board (Thorlabs, EDU-VS1/M), about 2.5 mm thick and a piece of 3mm-thick chicken breast as diffusers to mimic the scattering conditions. The optical path is sketched in Fig. S3. The laser beam at 532 nm was expanded and emitted onto a digital micromirror device (DMD, Texas Instruments, 0.65 inches, 1920×1080), which provided high rates of binary spatial light modulation. Handwritten digits and letters randomly selected from MNIST[3] and NIST[4] databases, respectively, were generated into 512×512 pixel patterns in the center of the DMD screen, to be consistent with the setup in Li *et al.*[1] to compare the reconstruction. The DMD generated patterns were projected on the PS board, transmitted through the diffuser, and acquired by a camera from the other side. To mimic the spatial inhomogeneity of scattering media and enlarge the robustness of our learning algorithm, we assembled the diffusers (PS board and chicken breast) on a translational stage to provide diverse scattering conditions, i.e., the scattering layer kept moving during data acquisition.

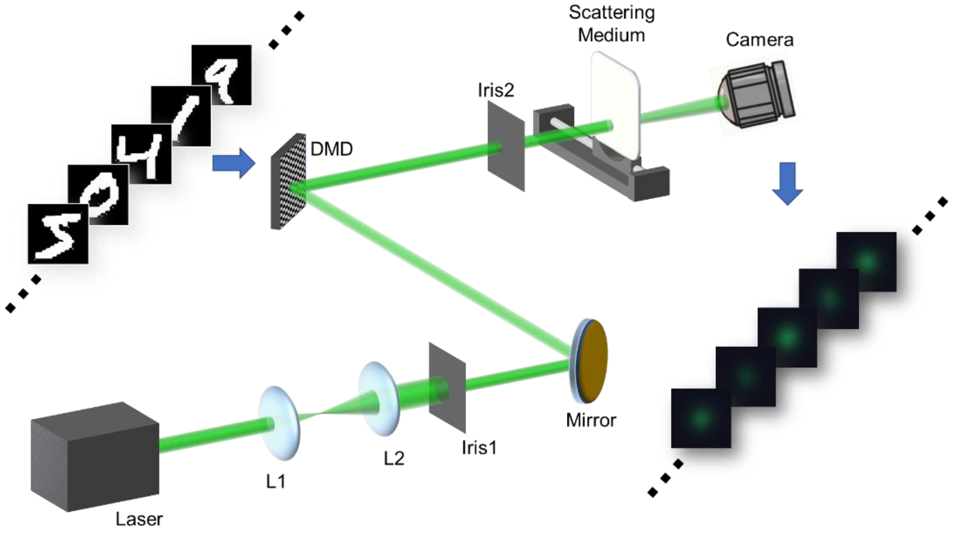

**Fig. S3** Schematic of the optical path on reconstructing images through a synthetic diffuser. L1, L2, lenses I1, I2, irises; DMD, digital micromirror device.

The digit and letter patterns are chosen from the MNIST and NIST datasets, respectively, which were used as the source and target domains for the DANN training, which adopted the cross-domain image-to-image translation of a limited number of unpaired data. Therefore, the DANN based network reconstructs the cross-domain patterns in the absence of "ground truths", which implies the feasibility of a similar network on reconstructing physiological vascular networks, when the paired "true" images are difficult to access. The classes branch and the domain branch, supervised by the classification labels and the domain labels, respectively, were connected to the middle layer of the UNet[5] through a Gradient Reversal Layer[2], which helped to indiscriminate the paired MNIST dataset and the unpaired NIST dataset with respect to the domain shift.(Fig S8) The data augmentation consisted of scaling, translational and rotational jitters that rendered the learning method more robust.

Apart from the domain adversarial training scheme used by Li *et al.*[1] for classification problem, our method used the DANN loss function for the mutual information disentanglement and validated the effectiveness of DANN for image-to-image transfer problem. . (Fig. S4) The training from the MNIST dataset, the source domain consisted of paired digits, was transferred to the NIST dataset, the target domain consisted of unpaired letters. Reconstruction results and comparisons are shown in Fig. S5 and S6, validating the feasibility of domain adaptation in imaging through scattering media.

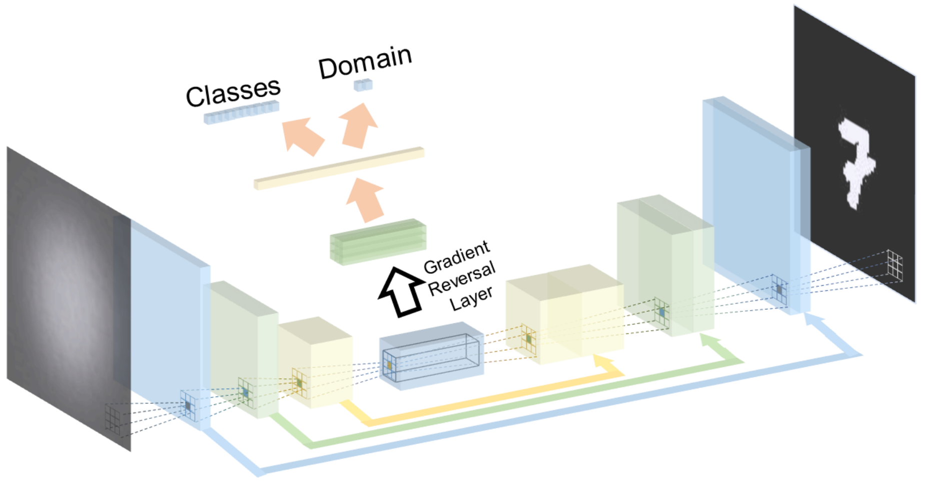


**Fig. S4** Structure of the neural network of our algorithm used in the *in vitro* experiment. It consisted of the UNet[5] as the backbone and two classification branches after a Gradient Reversal Layer that encouraged the emergence of indiscriminative features with respect to the classes label and domain source.

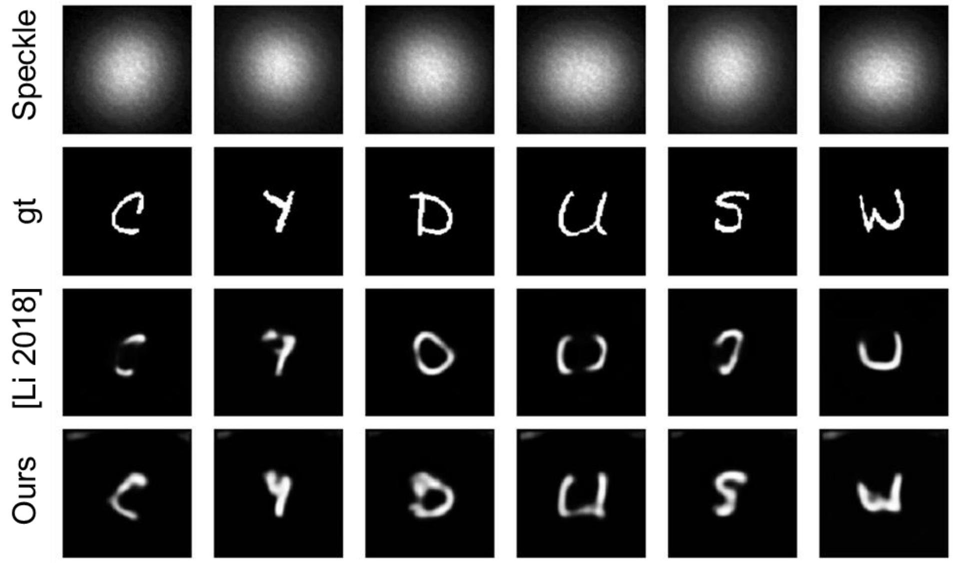


**Fig. S5** Comparison of the performance of Li *et al.* and our DANN-based algorithm using the **PS board** as a scattering medium. We trained both algorithms on handwritten digit dataset and performed testing on handwritten letter dataset. First row: speckle images captured by the camera. Second row: ground truth images sent to DMD for display. Third row: reconstructed patterns using the network of Li *et al.*. Fourth row: reconstructed patterns using our algorithm.

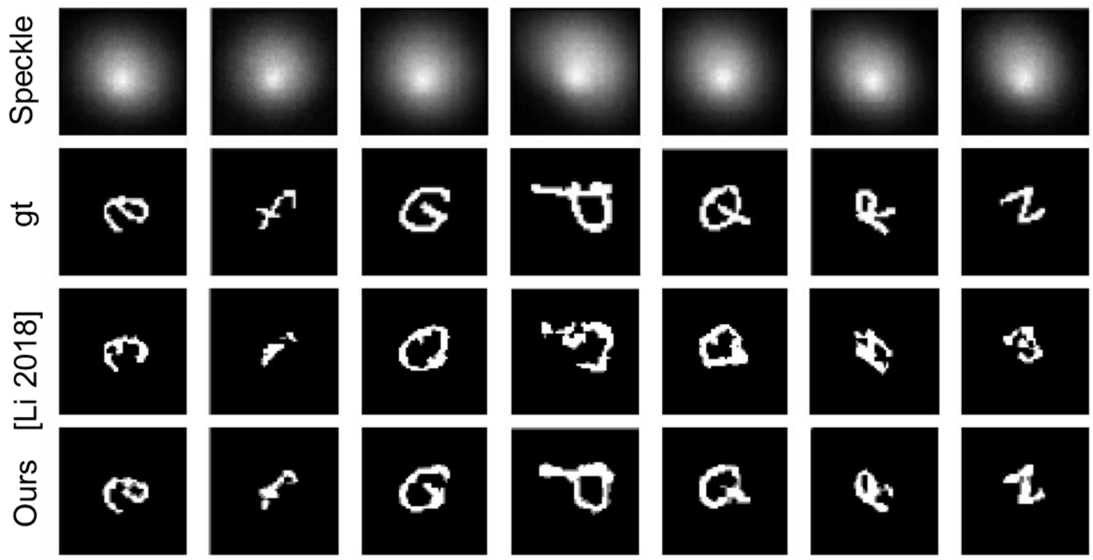


**Fig. S6** Comparison of the performance of Li *et al.* and our DANN-based algorithm using a piece of 3mm-thick chicken breast as a scattering medium. We trained both algorithms on handwritten digit dataset and performed testing on handwritten letter dataset.

### 4. Experimental setup for *in vivo* cerebral and bowel vascular imaging

Excitation light was provided by a 785nm laser source coupled to an 8 mm collimator. Light was expanded to illuminate the entire mouse head with the hair removed. The excitation power density at the imaging plane was about 29 mW·cm$^{-2}$ in our experiments, significantly lower than the reported safe exposure limit of 296 mW·cm$^{-2}$ at 785 nm. The emitted fluorescence was filtered by an 800-820 nm bandpass filter (Thorlabs, FBH810-10) and captured by a 4000*3000 pixel CMOS camera (Blackfly, BFS-U3-120S4M-CS) through a lens of focal length 25mm. We crop the central 2000*2000 square of camera pixels since it is enough to cover the whole range of interest. The camera was set above the mouse scalp to expose continuously in a frame rate of 25 frames per second, enabling recording the dynamic process. We record about 40 seconds (1000 images) for each mouse as soon as the injection of Indocyanine green begins. The sketch of creating thrombosis in a mouse cerebrum is shown in Fig. S7a. The optical configuration of fluorescence imaging is shown in Fig. S7b.

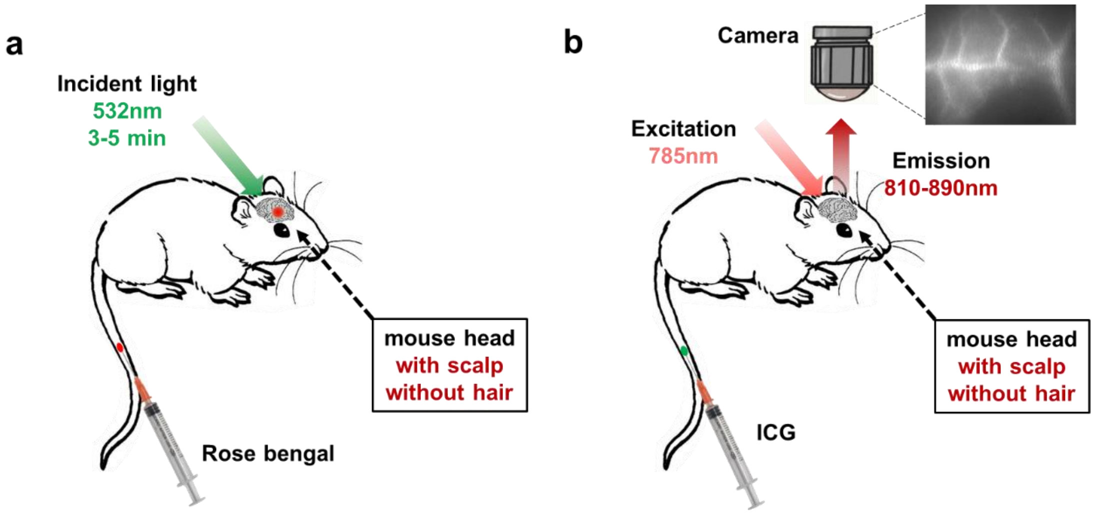


**Fig. S7** (a) Sketch of creating thrombosis in a mouse cerebrum. (b) Sketch of fluorescence imaging of a mouse cerebral vasculature.

The whole setup of imaging through mouse abdomens is the same as the aforementioned fluorescence brain imaging. After hair removal on the abdomen, the mouse was fixed on the imaging table, with the abdomen facing the camera. Since the area of mouse abdomens is larger than mouse heads, the central 2600*2600 square of camera pixels is cropped during the imaging process.

### 5. Validation and analysis of augmenting cerebral vasculature from different mice

References of cerebral vasculature were obtained by cutting off the scalp and taking the bright-field images of the mice cerebrum. We compared the multi-dimensional vasculature reconstruction in different mice to validate the correctness and stability of our VasNet algorithm.

We performed fluorescence imaging before and after thrombosis modeling on one **same** mouse. The mice were kept alive until the fluorescence images of the healthy cerebrum and the thrombosis cerebrum were acquired. The mice were then killed and cut-off the head skin to photo the cerebrum as the judgement for reconstruction.

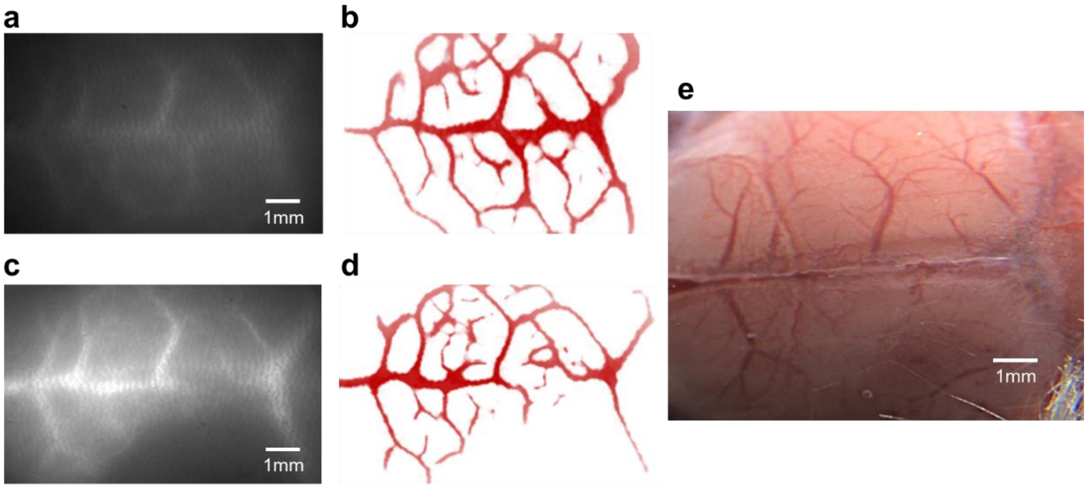


**Fig. S8** Comparison between inputs, outputs and the reference cerebral images of the mouse in the article. (a, c) Raw fluorescence images of the cerebral vasculature of a healthy mouse and a thrombosis mouse. (b, d) Outputs from the VasNet of the cerebral vasculature. (e) The bright-field reference image of the mouse cerebral vasculature with the scalp cut off.

Apart from this mouse, we also performed the same imaging experiments on another 2 healthy mice and 2 mice with thrombosis for validation. Note that these 4 healthy mice were different individuals.

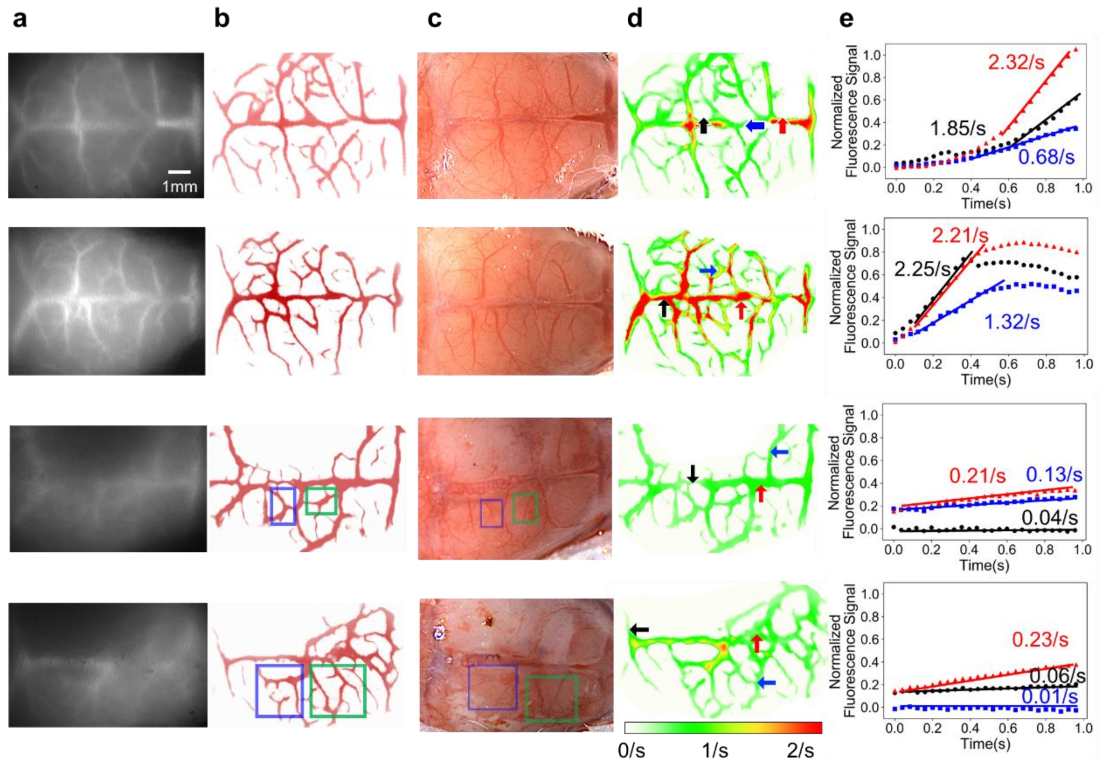


**Fig. S9** Comparison between inputs, outputs and the reference cerebral images of 2 healthy mice (the top two rows) and 2 thrombosis mice (the bottom two rows). (a) Raw fluorescence image of the cerebral vasculature through the scalp. (b) Output results of our VasNet algorithm corresponding to (a). (c) Bright-field reference images of the mice cerebral vasculature with the scalp off. (d) Normalized fluorescence signal distribution. € Normalized fluorescence intensity changes of the three positions in (d). The color bar in (d) representS the frequency of vessel illumination, initiated when the ICG was injected.

### 6. Validation of VasNet augmentation of vessels and bleedings in DSA images

Apart from the example shown in the article, here we provide another two examples of VasNet augmenting augmentation of vessels and bleedings in DSA images. In both figures, (a) is the original DSA imaging result shown on the hospital apparatus, and (b) are zoom-in time-sequenced details in the boxed area of (a), which all appear visually fuzzy. (c) and (d) are the outputs of our vasculature extraction algorithm and its zoom-in time-sequenced details respectively, which are explicit and can display the bleeding features apparently. (e) uses a color bar to show the evolution of bleedings according to time, where red represents the bleeding information which appears the earliest and blue the latest.

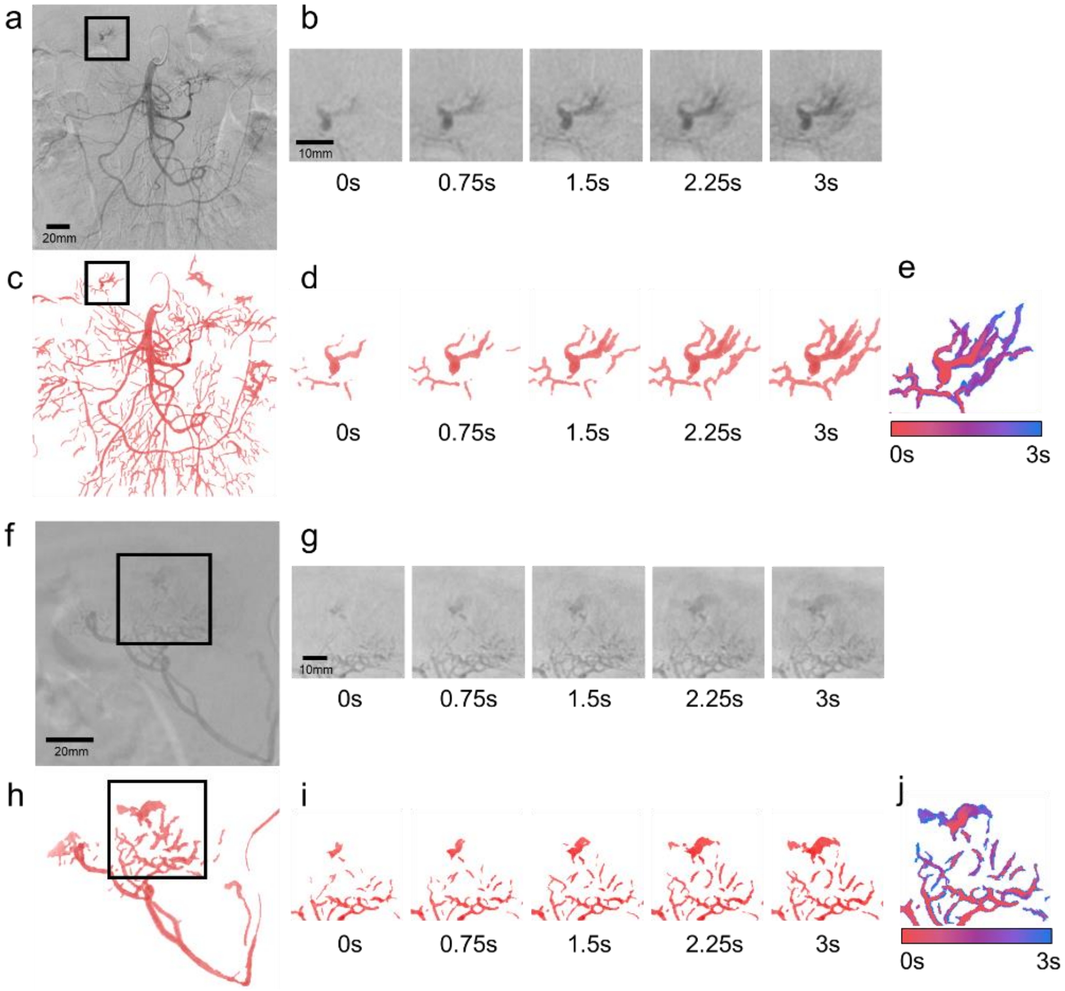


**Fig. S10** Two additional examples of augmenting vessels and bleedings in DSA images based on our algorithm.

### 7. Illustration of outliers in colitis classification statistics

The mice was grouped into Day 0, Day 2, Day 4, Day 6 to represent the number of days after being administrated with the DSS solution. Day 0 is the control group. Each group was composed of 5-6 mice with blinding. For each mouse, we performed fluorescence imaging through the abdomen and recorded the fluorescent process for 40 seconds. Since ICG molecules transport rapidly with circulation, only several tens of images contain meaningful information of the vasculature of guts. We randomly choose 3-10 meaningful images for each mouse, forming 51, 38, 34, 30 images for Day 0, Day 2, Day 4, Day 6 groups respectively.

We observe two prominent outliers in our data from Fig S6(a). The first one is a single point in Day 2 Group boxed by a green rectangle. After we performed dissection to this mouse, we observe clear irritation and swelling of its guts (Fig S6(b)), indicating this mouse has severe inflammation symptoms, though it belongs to Day 2 Group. The second one is the maximum point in Day 6 group boxed by a red rectangle, which belongs to a mouse with no dominate symptom of colitis, while all the other five mice in Day 6 Group have started hemafecia. The two outliers indicate the reliability of our statistical analysis.

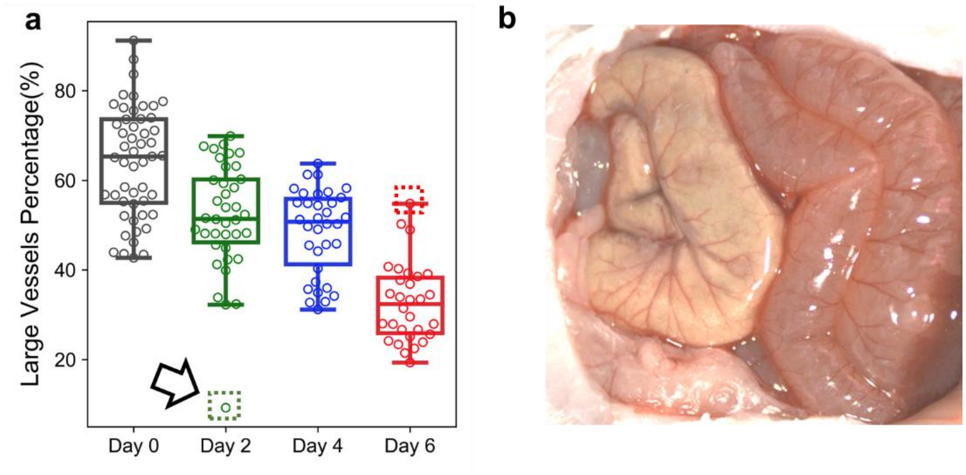


**Fig. S11** (a) Boxplot and swarm plot of the area percentage of large vessels. (b) The dissection image of the mouse corresponding to the single outlier in Day 2 Group, boxed with a green rectangle.
